# Spooky interaction at a distance in cave and surface dwelling electric fishes

**DOI:** 10.1101/747154

**Authors:** Eric S. Fortune, Nicole Andanar, Manu Madhav, Ravi Jayakumar, Noah J. Cowan, Maria Elina Bichuette, Daphne Soares

## Abstract

Glass knifefish (*Eigenmannia*) are a group of weakly electric fishes found throughout the Amazon basin. We made recordings of the electric fields of two populations of freely behaving *Eigenmannia* in their natural habitats: a troglobitic population of blind cavefish (*Eigenmannia vicentespelaea*) and a nearby epigean (surface) population (*Eigenmannia trilineata*). These recordings were made using a grid of electrodes to determine the movements of individual fish in relation to their electrosensory behaviors. The strengths of electric discharges in cavefish were larger than in surface fish, which may be a correlate of increased reliance on electrosensory perception and larger size. Both movement and social signals were found to affect the electrosensory signaling of individual *Eigenmannia*. Surface fish were recorded while feeding at night and did not show evidence of territoriality. In contrast, cavefish appeared to maintain territories. Surprisingly, we routinely found both surface and cavefish with sustained differences in electric field frequencies that were below 10 Hz despite being within close proximity of less than one meter. A half century of analysis of electrosocial interactions in laboratory tanks suggest that these small differences in electric field frequencies should have triggered the jamming avoidance response. Fish also showed significant interactions between their electric field frequencies and relative movements at large distances, over 1.5 meters, and at high differences in frequencies, often greater than 50 Hz. These interactions are likely envelope responses in which fish alter their EOD frequency in relation to changes in the depth of modulation of electrosocial signals.

## 1 INTRODUCTION

Gymnotiformes are a group of nocturnal fishes characterized by a suite of adaptations which allow them to localize conspecifics (Davis and Hopkins, 1988) and capture prey (Nelson and MacIver, 1999) in complete darkness. These fishes produce a weak electric field, typically less than 100 mV/cm (Assad et al., 1998), using an electric organ located along the sides of the animal and in its tail (Heiligenberg, 1991). This electric field, known as the electric organ discharge or EOD, is detected using specialized electroreceptors embedded in the skin. These receptors encode modulations generated through interactions with the electric fields of conspecifics and by nearby objects. This system provides a mechanism for communication among conspecifics and for the detection and characterization of prey and other salient environmental features (Nelson and MacIver, 1999; Crampton, 2019; Pedraja et al., 2018; Yu et al., 2019).

The nocturnal life histories of Gymnotiform species make them well suited for life in caves. Interestingly, a single species of Gymnotiform fish, *Eigenmannia vicentespelaea*, has been discovered in a cave system in central Brazil (Triques, 1996; Bichuette and Trajano, 2006, 2017). These fish exhibit features commonly found in species adapted to life in caves, including reduced pigmentation and reduction and/or elimination of the eyes (Culver and Pipan, 2019). The population is estimated to be around only 300 individuals (Bichuette and Trajano, 2015).

Fish in the genus *Eigenmannia* produce quasi-sinusoidal electric signals that are maintained at frequencies between about 200 and 700 Hz (Heiligenberg, 1991). Individual *Eigenmannia* change their electric field frequencies in response to electrosensory signals produced by nearby conspecifics (Watanabe and Takeda, 1963). In the Jamming Avoidance Response (JAR), individuals raise or lower their electric field frequency to avoid differences of less than about 10 Hz (Heiligenberg, 1991). Fish also change their electric field frequencies in relation to the relative movement between individuals that are encoded in the amplitude envelope of their summed electric fields (Stamper et al., 2013; Metzen and Chacron, 2013; Thomas et al., 2018; Huang and Chacron, 2016). Envelope responses can also occur in groups of three or more fish when the pairwise differences of their electric field frequencies are close, with 1-8 Hz, of one another (Stamper et al., 2012).

To discover the potential consequences of adaptation for troglobitic life on electrosensory behavior, we compared the electric behavior and movement of the cavefish *Eigenmannia vicentespelea* to nearby epigean (surface) relatives, *Eigenmannia trilineata*, that live in the same river system (Fig. 1A). We used a recently developed approach for characterizing electric behaviors and locomotor movements of weakly electric fishes in their natural habitats (Madhav et al., 2018; Henninger et al., 2020). This approach, which uses a grid of electrodes placed in the water, permits an estimation of the electric field of each fish and an analysis of their concurrent movement.

**Figure 1.**
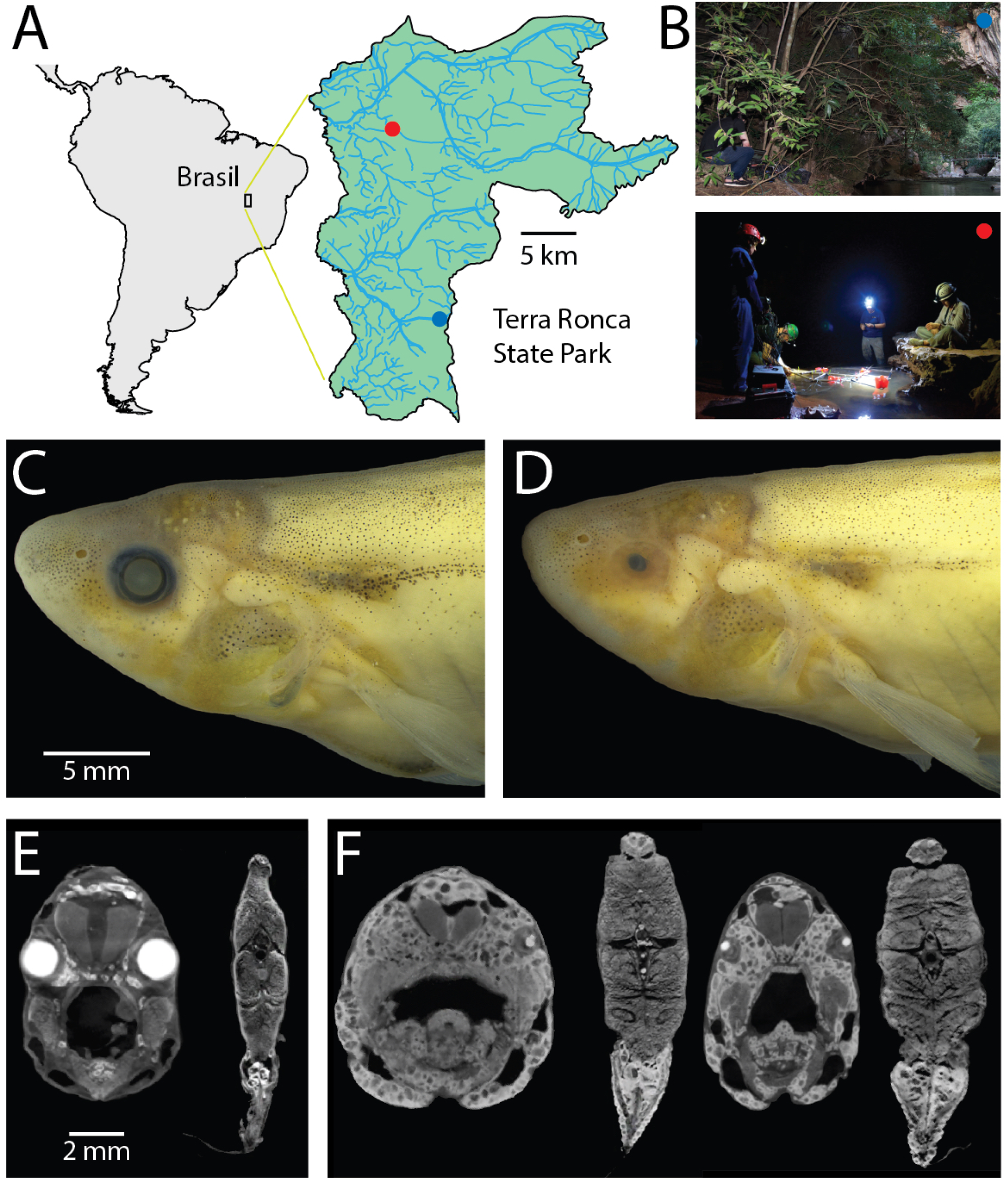
Surface and cave *Eigenmannia*. A) Study sites are in a clear water system in the Rio da Lapa karst region in Goiás, Brazil. B) Top, the surface *Eigenmannia* site was in the entrance of the Rio da Lapa cave. B) bottom, the cave *Eigenmannia* were found in the São Vicente II cave. C) Surface *Eigenmannia* have well-developed eyes and distinctive markings. D) Cave *Eigenmannia* have poorly developed or missing eyes and lack pigmented features. diceCT imaging reveals the differences in eye sizes and a potential difference in the relative size of electric organs. E) Coronal sections through the head and mid-body of a surface fish and F) two cavefish (dorsal is up). Large, bright cells in the caudoventral coronal sections appear relatively larger in the cave versus surface species.

## 2 METHODS

These observational studies were reviewed and approved by the Rutgers Institutional Animal Care and Use Committee and follow guidelines established by the National Research Council and the Society for Neuroscience.

### 2.1 Study sites

The study sites were located in Terra Ronca State Park (46°10’–46° 30’ S, 13° 30’–13° 50’ W), in the Upper Tocantins river basin, state of Goiás, central Brazil (Fig. 1A). We measured the electric behavior of the cavefish *Eigenmannia vicentespelaea* in the São Vicente II cave (13°58’37” S, 46°40’04” W) in October of 2016 (Fig. 1B). The electric behavior of the epigean species, *Eigenmannia trilineata*, was measured in the Rio da Lapa at the mouth of the Terra Ronca cave (13° 38’44” S; 46° 38’ 08” W) in April of 2016 (Fig. 1B). These streams have moderate water currents, clear water with conductivity below 20 *µ*S, and the substrate is composed of sand, rocks, and boulders.

### 2.2 Anatomy

Four alcohol fixed specimens from the collection at the Universidade Federal de São Carlos (Dr. Bichuette) were submerged in 11.25 Lugol’s iodine (I2KI) solution for up to 36 hours prior to diffusable iodine based contrast enhanced computer tomography (DiceCT). Stained specimens were removed from Lugol’s solution, rinsed in water to remove excess stain and sealed in rubber sleeves to prevent dehydration. Samples were then loaded into 50 mL polypropylene centrifuge tubes for scanning.

Stained and unstained specimens were scanned at the Core Imaging Facility of the American Museum of Natural History (New York, NY), using a 2010 GE Phoenix v-tome-x s 240CT high-resolution microfocus computed tomography system (General Electric, Fairfield, CT, USA). DiceCT scanning permits visualization of soft tissue details of the head and the body. Scans were made at 125 kV, with an exposure time of 60 seconds. Voxel sizes were 20.0-25.9*µ*m. Volume reconstruction of raw X-ray images were achieved using a GE Phoenix datos—x.

### 2.3 Recordings of electric behavior at field sites

*Eigenmannia* were initially identified and located using hand-held single-electrode probes with a custom amplifier/speaker system. This river system also permits direct visualization of the animals - the water is sufficiently clear and free of debris to see and photograph the fish (Hero Cam, GoPro 3, USA) when swimming in open water (see Supplemental videos S1, S2). Electric recordings were made using an array of amplified electrodes (50 cm between electrodes) (Madhav et al., 2018). For measurements of the epigean fish, an array of 8 electrodes was placed along the edges of the Rio da Lapa stream after sundown when the fish were active. In the São Vicente II cave, an array of 16 electrodes was placed in eddies and side pools along the primary stream. The flow at the center of the stream was too strong for grid array. Unfortunately, we were unable to use the larger grid on our second visit to the surface site due to a concurrent religious festival. As a result, the maximum XY range of the surface grid is about 100 cm diameter smaller than the cave grid. In all other respects, the measurements from each grid are identical.

We used an algorithm (Madhav et al., 2018) (code available at doi:10.7281/T1/XTSKOW) to identify each *Eigenmannia* using its time-varying fundamental frequency. We calculated an estimate of the strength of the electric field for each fish. Fish were considered to be ideal current dipoles—a source-sink pair of equal but time varying strength *I*(*t*), separated by a small distance *d*. The electric current dipole moment for the fish is defined as *p* = *Id* which has the units of Ampere-meter or “Am.” We calculated an estimate of the position and pose of each fish. Position and pose were calculated in relation to the distribution of power at each EOD frequency across the grid of electrodes. In these recordings, which were made in shallow water of no more than 40 cm depth with a planar array of electrodes, the position estimates were restricted to the XY-plane.

Continuous recording sessions using the grid were made both at the cave site (*N* = 14) and surface site (*N* = 5) of at least 600 seconds to over 1200 seconds duration. Intervals between recording sessions ranged from 5 minutes to several hours. Because fish could not be tracked between recording sessions, it is likely that some individual fish were measured across recording sessions.

### 2.4 Analysis of position and EOD frequency data

The position and EOD frequency data were analyzed, unless otherwise described, in 300 second epochs with 50% overlap. All analyses were conducted using custom scripts in Matlab (Mathworks, Natick MA). These data and scripts will be made freely available upon publication of this manuscript (URL). We estimated the XY region of movement for each fish for each epoch using a minimum convex polygon fitted to its positions. We then calculated the pairwise overlap between each convex polygon. Euclidean distances between pairs of fish were computed as a function of time during each 300 second epoch.

To assess the relations between pairs of EOD frequencies and their relative movement, we calculated Pearson correlations between 1) instantaneous distance between pairs of fish (distance) and 2) instantaneous difference in EOD frequency (dF). Pearson correlations were measured over 300 second epochs with 150 second overlap between epochs. These “dF/distance correlations” ranged from −0.93 to 0.90. The values for dF/distance Pearson correlations matched our impressions of the strengths of relations between dF and distance by visual inspection of the data. To estimate the rate of spontaneous correlations between dF and distance, we shuffled (Matlab randperm) the epochs of dF and distance measurements.

## 3 RESULTS

### 3.1 General observations

*Eigenmannia* at both the surface and cave sites were found in clear water streams with rock and sand substrates (Bichuette and Trajano, 2003, 2006, 2015, 2017). At the surface site, there was a marked diurnal modulation of behavior. During the day, fish were found alone or in groups along the banks of the Rio da Lapa, typically below or around boulders and rocks. The grid system was not used to make recordings of the fish during the day due to a local festival. Nevertheless, the distribution of *Eigenmannia* during the day appeared to be similar to that described previously at sites in Ecuador (Tan et al., 2005). Unlike in previous measurements at other study sites (Stamper et al., 2010), no other Gymnotiform species were detected in our short survey. At night, we observed *Eigenmannia* swimming in open water in the center of the stream, typically near the bottom.

Surface fish foraged in sandy substrates. Foraging behavior included hovering with slow forward or backward swimming punctuated by strikes into the substrate (see Supplemental video S1). These strikes involved tilting of the head and body downwards with a rapid forward lunge to drive the mouth into the sand a few millimeters. These strikes differ from feeding behavior described in *Apteronotus albifrons* in which fish captured freely swimming *Daphnia*, typically from below the prey (Nelson and MacIver, 1999). We observed groups of *Eigenmannia* simultaneously foraging with inter-fish distances on the order of 10s of centimeters. These distances suggest that fish experience significant ongoing electrosensory interference from conspecifics. At other locations, we visually observed single fish foraging while swimming rather than hovering a particular location.

At the cave site, fish did not exhibit diurnal modulation of behavior. Cavefish were observed along the banks of the stream and in small eddies and pools. Fish were alone or in small groups of up to 10 individuals. Cavefish retreated to crevices and spaces within boulders and rocks when disturbed, forming temporary aggregates of individuals (see Supplemental video S2). Videos show eyeless cavefish orienting face-to-face during social interactions. Such movements, in the absence of visual cues, must be controlled, at least in part, using electrosensory signals. The distinctive substrate foraging behavior routinely seen in the surface fish was not observed in these cavefish.

### 3.2 Morphology

Surface fish (*Eigenmannia trilineata*) had large eyes that were circumferential and of the same size (Fig. 1C,E) whereas the cavefish (*Eigenmannia vicentespelaea*) had eyes in various states of degeneration, from microphthalmic to completely absent (Fig. 1D,F). Preliminary review of scans of four fish showed potential differences in the size of the electric organ (Fig. 1E,F), but additional material will be necessary for quantitative analysis of electric organ structure and physiology.

A prior study found that surface fish are smaller than the cavefish (Bichuette and Trajano, 2006): mean length of the snout to the end of the anal fin base was reported to be 8.45 cm (sd=2.67) in surface fish and 11.1 cm (sd=2.47) in cavefish. Size is important as it likely impacts the strength of electric fields: larger fish typically can generate larger currents in their electric organs.

### 3.3 Electric field amplitudes

The grid recording system permits an estimation of the strength of each individual’s electric field using Fourier analysis. The strength of electric fields (Ampere-meter, “A-m”) of surface fish were significantly lower than cavefish (Wilcoxon sign-rank, two-sided, z=2.98, p=0.0029) with mean strength of 6.5 *×* 10^−4^ A- m for surface fish and 9.66 *×* 10^−4^ A-m) for cavefish (Fig. 3B). The surface fish are generally shorter than the cavefish *Eigenmannia* and also appear to have relatively smaller electric organs. The increase in EOD amplitude may be an energetically costly adaptation (Markham et al., 2016) to life in this cave: larger EOD amplitudes would increase the ability to detect objects and capture prey (Nelson and MacIver, 1999) in the absence of visual cues.

### 3.4 Electric field frequencies

We used the grid recording system to calculate the EOD frequency of each fish within the grid (Fig. 2). EOD frequencies of surface fish were between 299.9 Hz and 435.6 Hz (N=110) whereas EOD frequencies of cavefish were between 230.0 Hz and 478.6 Hz (N=82) (Fig. 3A). The distribution of EOD frequencies of surface and cavefish are significantly different (Wilcoxon sign-rank, two-sided, z=2.57, p=0.0100). The mean and median frequencies of surface fish were 375.8 Hz and 382.0 Hz, and for cavefish were 356.6 Hz and 360.8 Hz. There was a dramatic diurnal modulation of behavior in surface fish, but grid data were collected only at night. The electric fields of some species of Gymnotiform fishes have been shown to have diurnal modulations and amplitude and frequency content (Stoddard et al., 2006; Markham et al., 2009; Migliaro et al., 2018; Sinnett and Markham, 2015). We have not observed diurnal modulation of EOD frequencies of *Eigenmannia* in previous field studies (Tan et al., 2005; Stamper et al., 2010).

**Figure 2.**
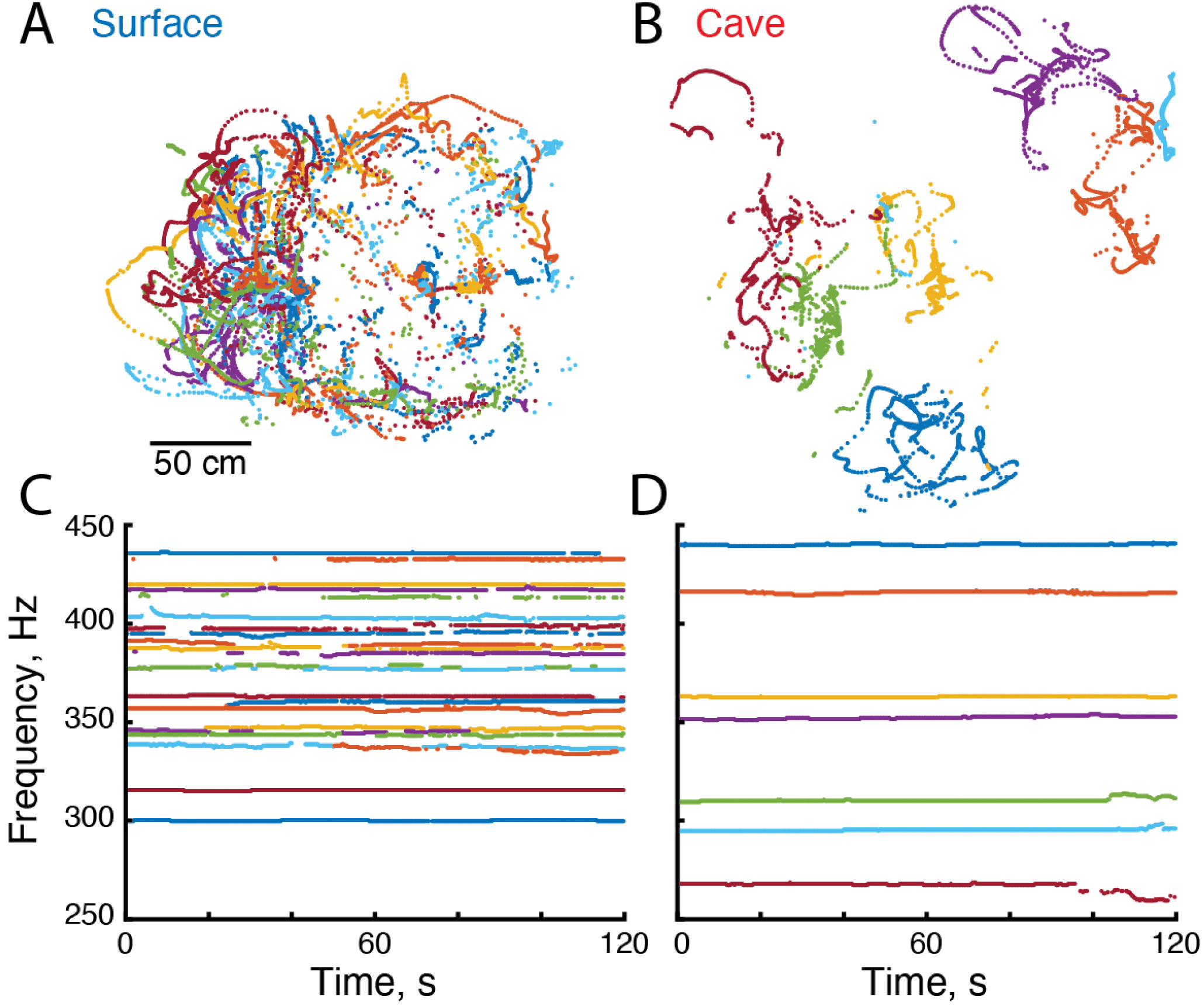
Electrical and locomotor behavior of *Eigenmannia*. A-B) Positions of surface and cave *Eigenmannia* over a period of 120 seconds. C-D) EOD frequencies of these fish. Each color represents a unique fish in each recording.

**Figure 3.**
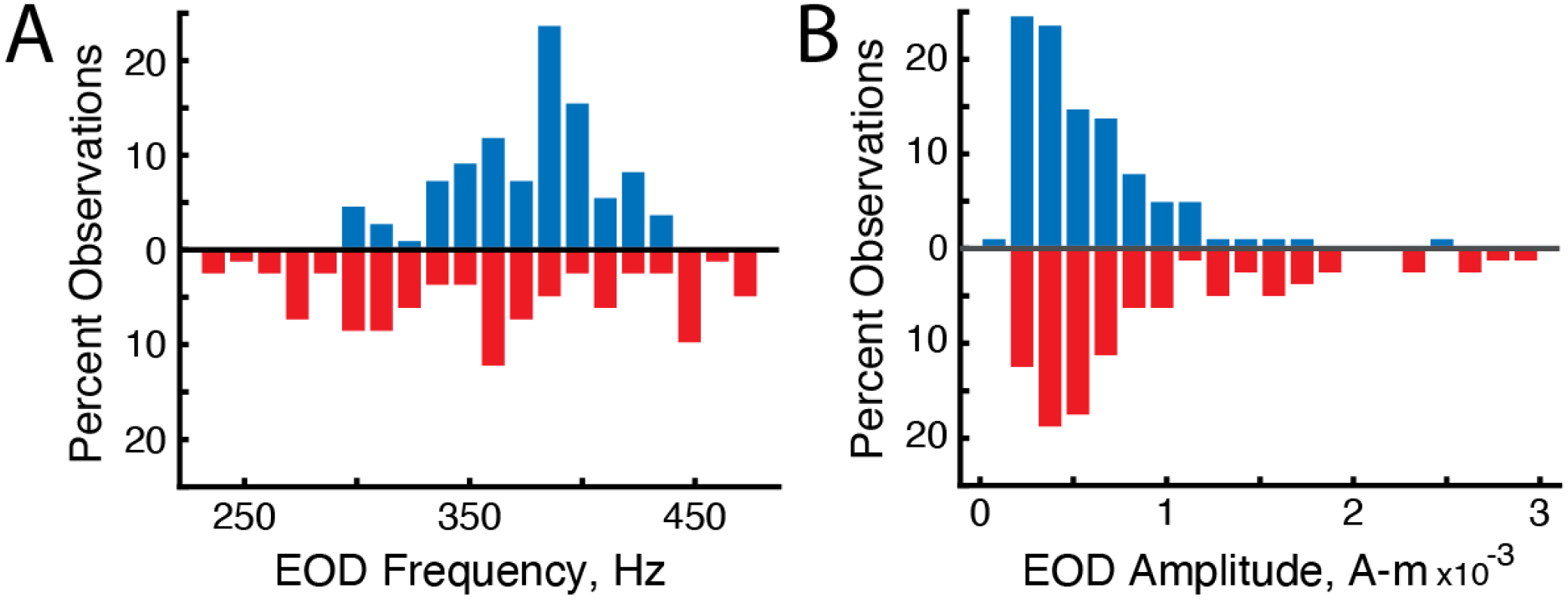
Differences in the distribution of EOD frequencies and amplitudes in surface fish (blue, up) and cavefish (red, down). A) Percent observations of EOD frequencies for surface fish and cavefish. B) Distribution of EOD amplitudes.

We observed variations in EOD frequencies that likely include social signals such as chirps and envelope responses (Fig. 2C,D). The variance of EOD frequencies in freely swimming groups of surface fish (mean=0.64 Hz, n=184) and cavefish (mean=1.10 Hz, n=292) were significantly different (T-test, p*<*0.0001, df=467, tstat=6.1). We expect the variance of EOD frequencies in surface fish will differ during daylight hours, when the fish are hiding along the shores of the rivers, in relation to night hours when the fish are active. Additional recordings made during the day are needed.

### 3.5 Differences in EOD frequencies

The JAR is a behavior that, in laboratory settings, reduces the likelihood that pairs fish will have a difference in EOD frequency (dF) of less than 10 Hz (Heiligenberg, 1991). In surface fish the mean dF was 40.32 Hz (sd=26.90, N=2251) while the cavefish the mean dF was 78.38 Hz (sd=47.81, N=3908) (Fig. 4A). The distribution of mean dFs was significantly larger in cavefish than surface fish (T-test p*<*0.0001, df=6157, tstat=34.74). These findings are consistent with a role of the JAR in maintaining larger dFs between most individuals, but may also simply reflect the greater density of fish that we observed at the surface site.

**Figure 4.**
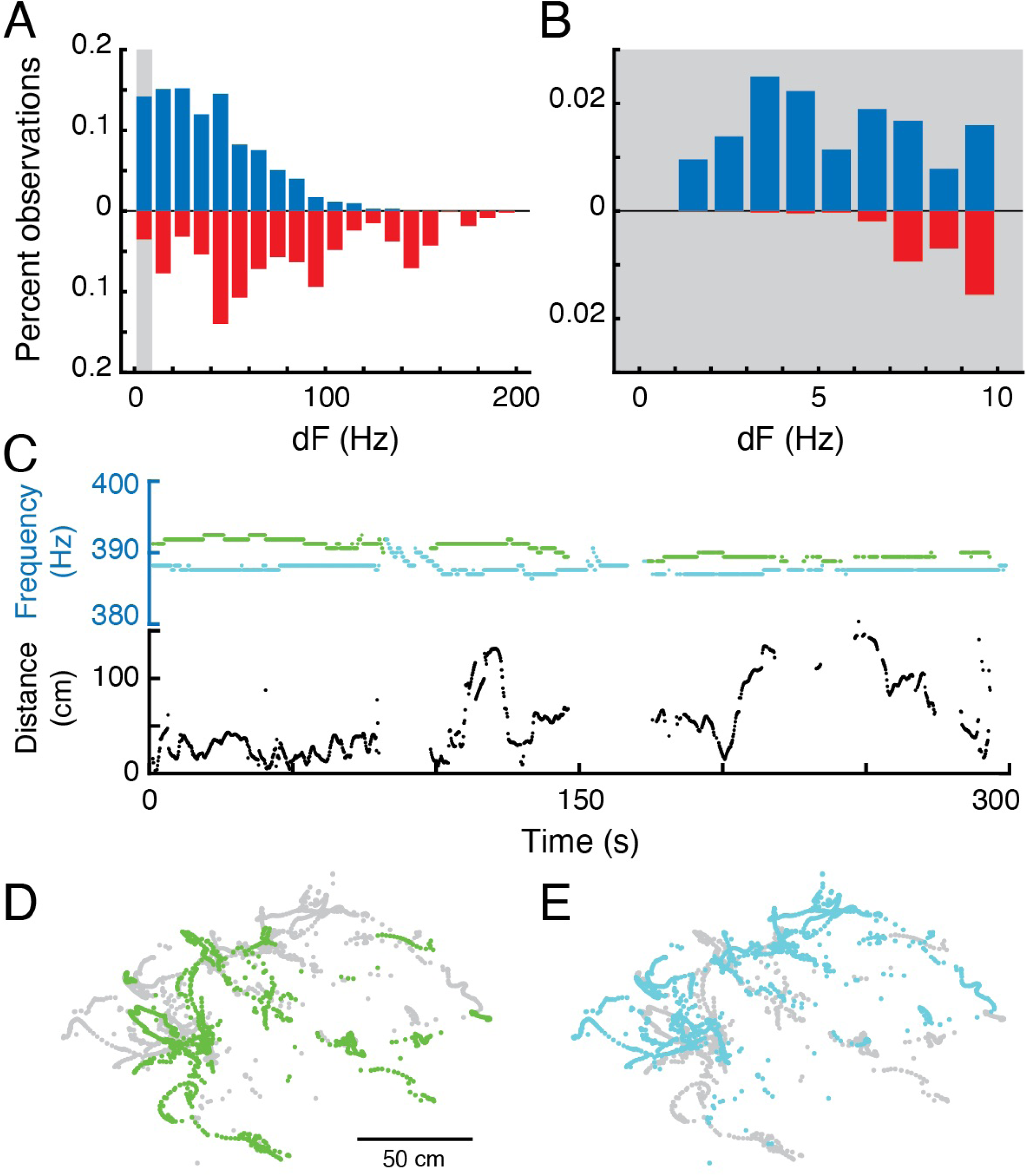
Differences in EOD frequencies. A) Percent observations of dFs for surface fish (blue, up) and cavefish (red, down). Grey bar highlights the low frequency region shown in B. C) Example epoch of two fish with sustained low-frequency dF. Top, EOD frequencies each fish; bottom Euclidian distance over time. D-E) X-Y positions of each fish.

Interestingly, we routinely found nearby fish with sustained, lasting for hundreds of seconds, differences in electric field frequencies that were below 10 Hz (Fig. 4B). In surface fish, there were 294 pairs with mean dFs of less than 10 Hz over 300 second duration epochs, and of those, 142 were less than 5 Hz (out of total 1947 pairwise interactions). In cavefish, there were 18 epochs with mean dFs that were less than 10 Hz, of which 4 were less than 5 Hz (out of total 435 pairwise interactions). Low dFs between pairs of fish were at distances within 50 cm (Fig. 4C-E).

### 3.6 Similarities and differences in movement patterns between surface and cavefish

During daylight hours, surface fish were found deep within crevices and other refugia along the sides of the Rio da Lapa, but emerged after dark for feeding and social behavior that occurred in open water throughout the stream. In contrast, cavefish were found swimming with no apparent variation with respect to time of day along the sides of streams and in pools and eddies.

We estimated the position of each fish in and around the grid over time (Madhav et al., 2018). Cavefish appeared to swim within small regions or territories on the order of tens of centimeters in diameter. In contrast, surface fish appeared to have widely overlapping swimming trajectories. To examine the relative movements of fish, we divided the data into 300 second duration epochs and fitted a minimum convex polygon to each fish’s positions.

Although there was no difference in the overall size of the convex polygons between surface and cavefish (T-test, p=0.5840, df= 499, tstat=0.5480), we found significantly more overlap in the trajectories of surface fish. The overlap between convex polygons of pairs surface fish (mean 13.89% overlap, std=9.57, n=1371) was significantly greater (T-test, p*<*0.0001, df=2219, tstat=13.22) than cavefish (mean 8.41%, std 9.08, n=950). Our impression is that the cavefish are territorial whereas surface fish were not territorial while feeding. The increased overlap in trajectories in surface fish may be a result of their lower amplitude electric fields, which could reduce the distance for detection of conspecifics, or simply due to the larger numbers of surface fish at the study sites. Further, we expect that the surface fish may defend territories during the day when hiding in refugia. The necessary grid recordings were not possible during the day—additional recordings are needed to better understand the diurnal modulation of social behavior in the surface fish.

### 3.7 Social stimuli and movement interact to modulate EOD signaling

Movement affected social signaling. Three surface fish were placed in tubes that restricted their movement during grid recordings. The variation of EOD frequencies in these fish (mean=0.39 Hz, std=0.36, n=29) was significantly lower than in freely swimming fish (T-test, p=0.0013, df=211, tstat=3.26). This is particularly interesting because the fish in tubes were surrounded by freely moving fish. These fish were exposed to conspecific electric fields that varied in amplitude as the other fish swam around them, producing envelopes (Stamper et al., 2013; Thomas et al., 2018). The dramatic reduction in the variation of their electric fields suggests that fish may modulate their electric signaling to conspecifics in relation to their own movement. Alternatively, other ancillary factors may have diminished overall sensorimotor responsiveness in restrained fish.

We also measured the EODs of three solitary cavefish—individuals that were freely moving but nevertheless isolated from conspecific electric fields. These fish also exhibited reduced variability of their EOD frequencies (mean=0.24 Hz, std=0.09, N=9) compared to cavefish in groups (T-test, p=0.0080, df=292, tstat=0.94). This variability of the electric field frequencies of solitary cavefish was not significantly different from the variability of surface fish in tubes (T-test, p=0.2554, df=36, tstat=1.16). These data are consistent with the theory that variability in EOD frequencies is driven by a combination of a fish’s own movement and its electrical interactions with conspecifics.

### 3.8 Correlations between movement and EOD frequencies suggest envelope responses in the field

We examined the relations between relative movement and EOD frequency as fish interacted with conspecifics. We measured the dFs of all pairs of fish and their simultaneous pairwise distances (Fig. 5). In many pairs of fish, dF and distance appeared to be correlated—either changing in the same or opposite directions (Fig. 5A,B). Negative correlations (e.g. dF increases as distance decreases) may be driven by a JAR-like response known as ‘envelope’ responses (Stamper et al., 2012, 2013; Metzen and Chacron, 2013; Thomas et al., 2018; Huang and Chacron, 2016). As *Eigenmannia* move closer together, the relative amplitudes of each fish’s electric field increases (similar to moving closer to a sound source). Fish respond to these increases by shifting their electric field frequency either up or down (Stamper et al., 2013; Metzen and Chacron, 2013; Thomas et al., 2018; Huang and Chacron, 2016).

**Figure 5.**
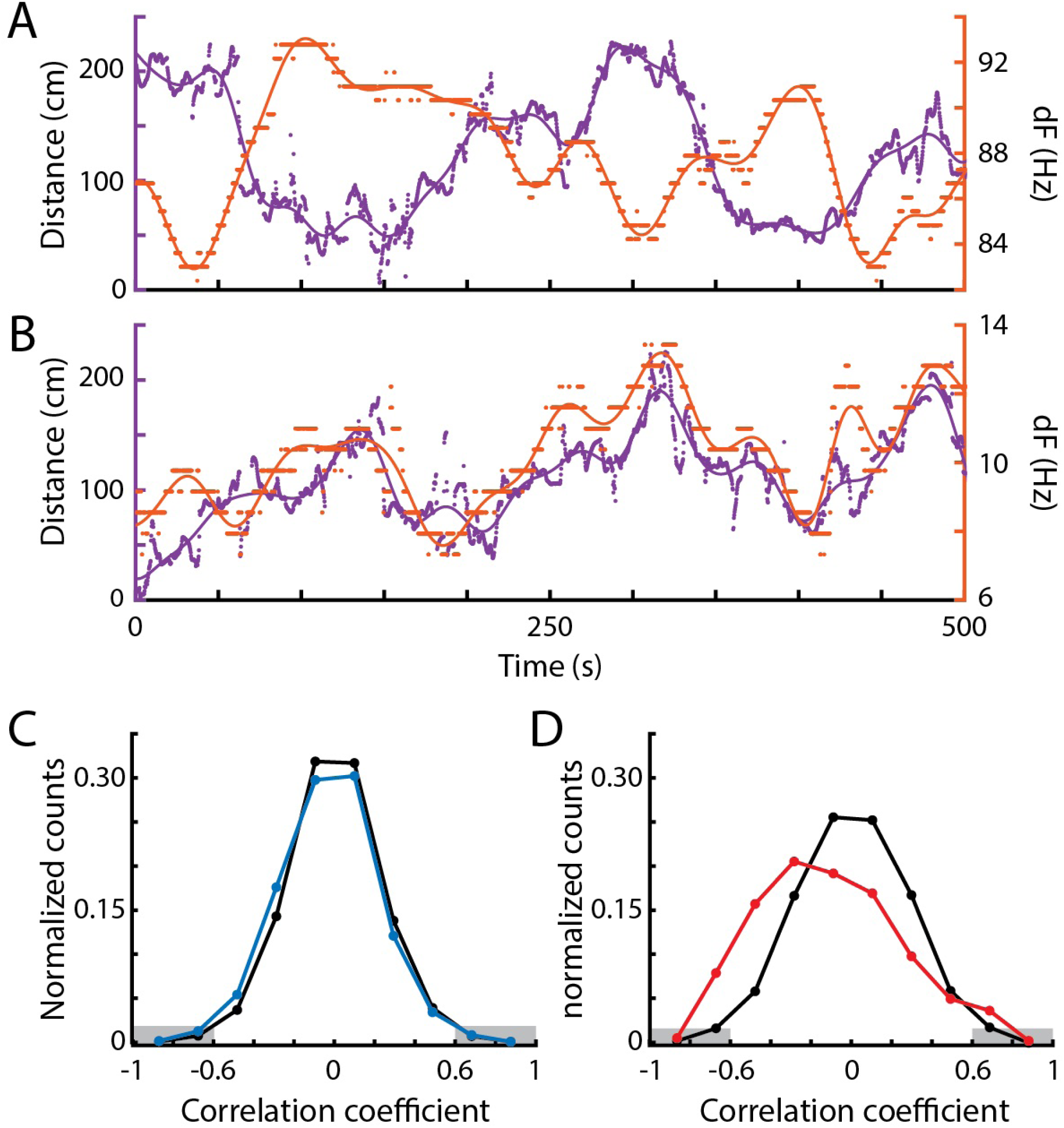
Correlations between EOD frequency and movement. A) Example of a strong negative correlation between distance (purple, left) and dF (orange, right) of a pair of *Eigenmannia* over a period of 5 minutes. Dots are measurements, lines are low-pass fits of the data. B) Example of a strong positive correlation. C) Distribution of Pearson correlations between distance and dF for surface fish (blue) and shuffled data (black). D) Distribution of Pearson correlations between distance and dF for cavefish (red) and shuffled data (black). There were significantly more strong negative Pearson correlations, below −0.6 (marked in grey) in both surface and cavefish. Unlike in surface fish, there were significantly more examples of high Pearson correlations (above 0.6) in cavefish than in shuffled data.

Correlations between two independently varying measurements may, however, occur spontaneously. To assess the rate of spontaneous correlations, we shuffled the distance and dF trajectories in time. We compared distributions of Pearson correlations between dF and distance in the shuffled and original data (Fig. 5C,D).

In surface fish, we found significantly more negative correlations but not positive correlations (Fisher’s exact test, less than 0.6: p=0.048, greater than 0.6: p=0.719)(Fig. 5C). In cavefish, we found both significantly more negative and positive correlations (Fisher’s exact test, less than 0.6: p*<*0.0001, greater than 0.6: p*<*0.0001)(Fig. 5D). These results suggest that both surface and cavefish exhibit JAR-like envelope responses—negative correlations between distance and dF. However, cavefish routinely exhibited positive dF-distance correlations, which may be produced during aggressive behaviors, such as active jamming (Tallarovic and Zakon, 2002).

Positive correlations were observed in fish with dFs of lower than 10 Hz (Fig. 5B). Such positive correlations at low dFs are unexpected because of the impact of the JAR. The JAR is strongest at dFs of approximately 2-8 Hz (Heiligenberg, 1991) and should increase dFs in nearby fish, generally resulting in negative correlations between distance and dF. These unexpected positive correlations may be a result of the production of social signals that ‘override’ the JAR, or may be driven by social interactions with other nearby fish that have higher dFs.

Interestingly, we found examples of strong Pearson correlations between dF and distance at dFs over 50 Hz distances of over 100 cm and (Fig. 5A). This is interesting because these distances were previously believed to be beyond the boundary of the animal’s ability to detect such signals (Nelson et al., 1997; Bastian, 1981; Henninger et al., 2018). These data are similar to evidence in other species of weakly electric fish which show that individuals can detect each other via weak modulations of their electric signals (Henninger et al., 2018; Raab et al., 2019).

## 4 DISCUSSION

We compared the electric behavior and locomotor movement in a population of epigean fish *Eigenmannia trilineata* and a population of troglobitic fish *Eigemannia vicentespelea*. We observed that surface fish had diurnal changes in behavior, hiding in refugia along the sides of streams during the day and swimming and feeding along the bottom of sandy and rocky streams during the night. In contrast, cavefish showed no apparent diurnal modulation of behavior.

### 4.1 Energetics

The EOD amplitudes of the troglobitic *Eigenmannia* were, on average, higher than the nearby surface fish. This may be in part due to body size: a previous report (Bichuette and Trajano, 2006) showed that the cavefish are generally larger than the surface fish. Further, our preliminary anatomical evidence from diceCT scans suggest that the electric organs of cavefish may be relatively larger than in the surface fish. Irrespective of size, the energetic cost of generating electric fields is high, consuming up to one quarter of an individuals energy budget (Salazar et al., 2013; Markham et al., 2016). The fact that cave *Eigenmannia* produce such energetically costly electric fields suggests that sufficient food resources are available and accessible. The loss of eyes and pigment in these cavefish, therefore, is likely not under strong selection for energetic costs, but rather neutral selection (Jeffery, 2009).

Indeed, Gymnotiform fishes throughout the Amazon basin have relatively small eyes. Over the years, we have encountered many individual fish with missing or damaged eyes. Further, we routinely observe dense infestations of nematode parasites in the eyes of individuals from a related genus, *Apteronotus*. These anecdotal observations are consistent with the theory that Gymnotiform fishes rely more heavily on electric sensing than vision for survival and reproduction. Gymnotiform fishes, including the troglobitic weakly electric cavefish *Eigenmannia vicentespelea*, represent a unique opportunity to study evolutionary changes related sensory perception and behavioral control.

### 4.2 EOD frequencies

The distribution of EOD frequencies in both of these groups of *Eigenmannia* were uni-modal, as has been previously observed (Tan et al., 2005). Further, we were not able to identify any frequency-dependent signalling that might be correlated with sex. Sex differences in EOD frequencies are well known in *Apteronotus* (Fugère et al., 2010; Raab et al., 2019) and *Sternopygus* (Hopkins, 1972), and there may be sex differences in EOD frequencies in *Eigenmannia* (Dunlap and Zakon, 1998). There was a difference in the distributions of EOD frequencies between the cavefish and surface fish, but the meaning of these differences remains unclear. The larger range of frequencies seen in the cavefish may be due to sustained interactions related to territoriality—fish in adjacent territories may increase their dFs over time.

Unexpectedly, we observed sustained, low-frequency dFs in both nearby surface and cavefish. Such interactions are not captured by models of the JAR (Madhav et al., 2013). Low-frequency dFs potentially impair electrolocation in *Eigenmannia* (Ramcharitar et al., 2005), and can be avoided by performing a JAR (Heiligenberg, 1991) or moving further apart (Tan et al., 2005). It seems unlikely that these fish were experiencing significant impairment of sensory function, as they were engaged in feeding on prey under the substrate in complete darkness. Studies in *Apteronotus albifrons* demonstrates that fish may use ampullary receptors to detect exogenously generated electric fields for prey capture (Nelson and MacIver, 1999). In addition, elasmobranchs have also been shown to detect and discriminate signals from substrate-bound prey using passive electroreception mediated by ampullary receptors (Kalmijn, 1971, 1982).

### 4.3 Territoriality

Territoriality is a form of space-related dominance (Kaufmann, 1983). The most prominent function of having a territory is to provide the holder with an secured supply of resources. In the epigean streams outside of the Terra Ronca cave, food resources for *Eigenmannia* appear to be widely distributed in sandy substrates. Our guess is that the size and distribution of prey items precludes territorial defense of food resources. On the other hand, we expect to find evidence of territoriality during the day, as refugia likely vary in quality and are relatively small.

Why cavefish exhibit evidence of territoriality is unclear. There are no known predators of *Eigenmannia* in the cave, eliminating the value of protective refugia. Territoriality may occur as a result of uneven distribution of food resources in the cave, due to other physical features that impact the fish, or a consequence of plesiomorphic social and/or reproductive behaviors. Territoriality has been described in other genera, such as *Gymnotus* (Zubizarreta et al., 2020).

### 4.4 Spooky interactions at a distance

The strength of electric fields in water decay at a rate of approximately distance cubed (Henninger et al., 2018). As a result, the distances between fish determine the strength of the interaction of their electric fields: nearby fish will experience higher EOD voltages than from those of distant fish. Because the EODs of *Eigenmannia* are nearly sinusoidal, distance will have an effect on the amplitude of modulations caused by the summation of EODs: nearby fish will have large amplitude modulations near 100%, whereas distant fish will have far lower depths of modulation, often below 10%. The relative movement between fish will cause concomitant changes in the strengths of EODs and depths of modulation that are proportional to distance. The changes are known as ‘envelopes’—the modulation of amplitude modulations. Envelope stimuli can elicit changes in electric field frequencies in *Eigenmannia* and other Gymnotiform species (Stamper et al., 2013; Thomas et al., 2018).

We found strong correlations between distance and pairwise differences in EOD frequencies at large distances of over 1.5 meters and dFs of over 50 Hz in both surface and cavefish. These results are similar to reports from field studies that examined other Gymnotiform species (Henninger et al., 2018; Raab et al., 2019). These field studies suggest that electric fish are far more sensitive to electrosocial stimuli than previously appreciated, requiring a reexamination of the neural systems for their perception.

Although many of the correlations between distance and dF were spontaneous, both surface and cavefish exhibited significantly more negative correlations between distance and dF than random. These negative correlations are likely driven by the amplitude envelope of electrical interference patterns, producing JAR-like behavioral responses (Stamper et al., 2012). These findings show that laboratory studies of envelope responses are ecologically relevant (Stamper et al., 2012, 2013; Metzen and Chacron, 2013; Thomas et al., 2018; Huang and Chacron, 2016). Cavefish also exhibited significantly more positive correlations between distance and dF. We believe that these may be aggressive signals in which fish actively jam each other (Tallarovic and Zakon, 2002).

#### 4.4.1 Permission to Reuse and Copyright

Figures, tables, and images will be published under a Creative Commons CC-BY licence and permission must be obtained for use of copyrighted material from other sources (including re-published/adapted/modified/partial figures and images from the internet). It is the responsibility of the authors to acquire the licenses, to follow any citation instructions requested by third-party rights holders, and cover any supplementary charges.

## AUTHOR CONTRIBUTIONS

Bichuette, Soares, and Fortune conceived and executed the field research. Andanar, Madhav, Jayakumar, and Fortune analyzed the electrical data. Soares collected and analyzed the diceCT data. Madhav, Jayakumar, and Cowan contributed to the design, use, and analysis of data from the grid system. All authors contributed to the preparation of the manuscript.

## ACKNOWLEDGMENTS

Field research permits were granted by the ICMBio and SEMARH/SECIMA. We would like to thank DF Torres, MJ Rosendo and CCP de Paula for help in the field. Thanks to Dr. Jessica Ware for the use of the CT scanner. This manuscript has been released as a pre-print at [https://www.biorxiv.org/content/10.1101/747154v1], (Fortune et al., 2019).

## SUPPLEMENTAL DATA

Supplementary Material should be uploaded separately on submission, if there are Supplementary Figures, please include the caption in the same file as the figure. LaTeX Supplementary Material templates can be found in the Frontiers LaTeX folder.

## DATA AVAILABILITY STATEMENT

The data and the Matlab scripts used to generate the figures will be made available on publication of this study on github (URL).

